## Supplemental Document 3 for "Genes of the Pig, *Sus scrofa*, reconstructed with EvidentialGene"

Supplemental Genome map pictures of Pig Genes, from Evigene, NCBI and Ensembl gene sets of June 2018. These display examples of Evigene improved models, with greater homology to human reference genes than Ensembl and NCBI, but for noted cases where NCBI model is equivalent or better in human gene alignment. Pig gene model comparisons are from Suppl. Table 2. Genome maps are from http://eugenes.org/EvidentialGene/vertebrates/pig/pig18evigene/map/

**Suppl. Figure 1, pigchr9nobox. Gene Name:** homeobox protein NOBOX; **Notes:** evg model has 2 more coding exons for greater human alignment, all have 1 model here, ncbi = ensembl; simple gene example, 8 vs 6 coding exons; human:NP_001073882.3, 691aa, Pig models: Evigene:Susscr4EVm006479t1, 574 aln, NCBI:NP_001182045.1, 267 aln, Ensembl: ENSSSCP00000027616.1, 267 aln

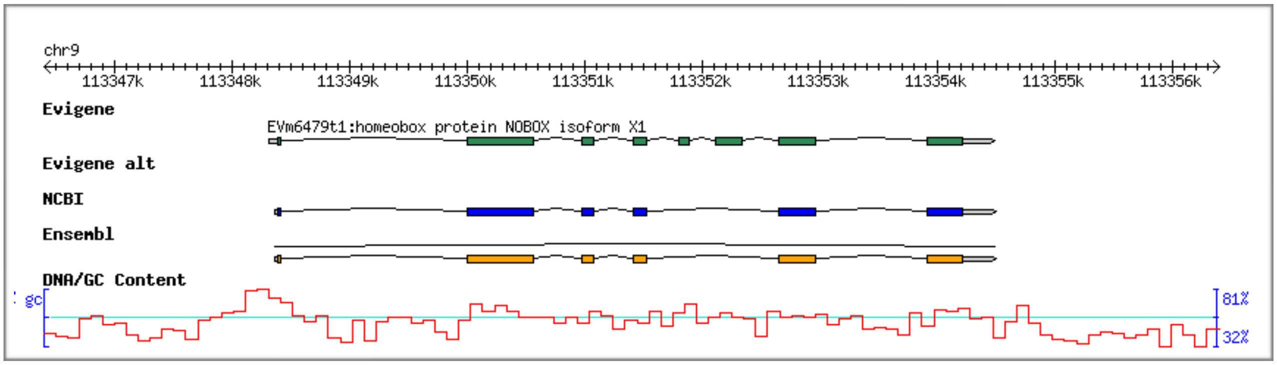


**Key to Gene maps**: Each picture is a genome map view of Pig gene models from Evigene, NCBI and Ensembl, all of 2018 June vintage, mapped onto chromosome set listed in Data citaions (version Sscrofa11.1 of 2018). Tracks (or rows) from top to bottom are (1) chromosome ruler in kilobases; (2) Evigene main/longest model; (3) Evigene alternate isoforms; (4) NCBI models (main, alternates and ncRNA); (5) Ensembl models (main, alternates and ncRNA); and (6) DNA/GC content percentage graph (blue line near 50% is average, 0% is gap).

**Suppl. Figure 2, pigchr5eeantigen.** **Gene Name**: early endosome antigen 1; **Notes:** evg many alts, ncbi 1 model, ensembl is missing this gene; Picture has cropped half of Evigene alts. Models: human:XP_011537116.1, 1453 aa, Pig Evigene:Susscr4EVm000897t1, 1485 align, NCBI:XP_020946919.1, 1470 align; Ensembl: missing

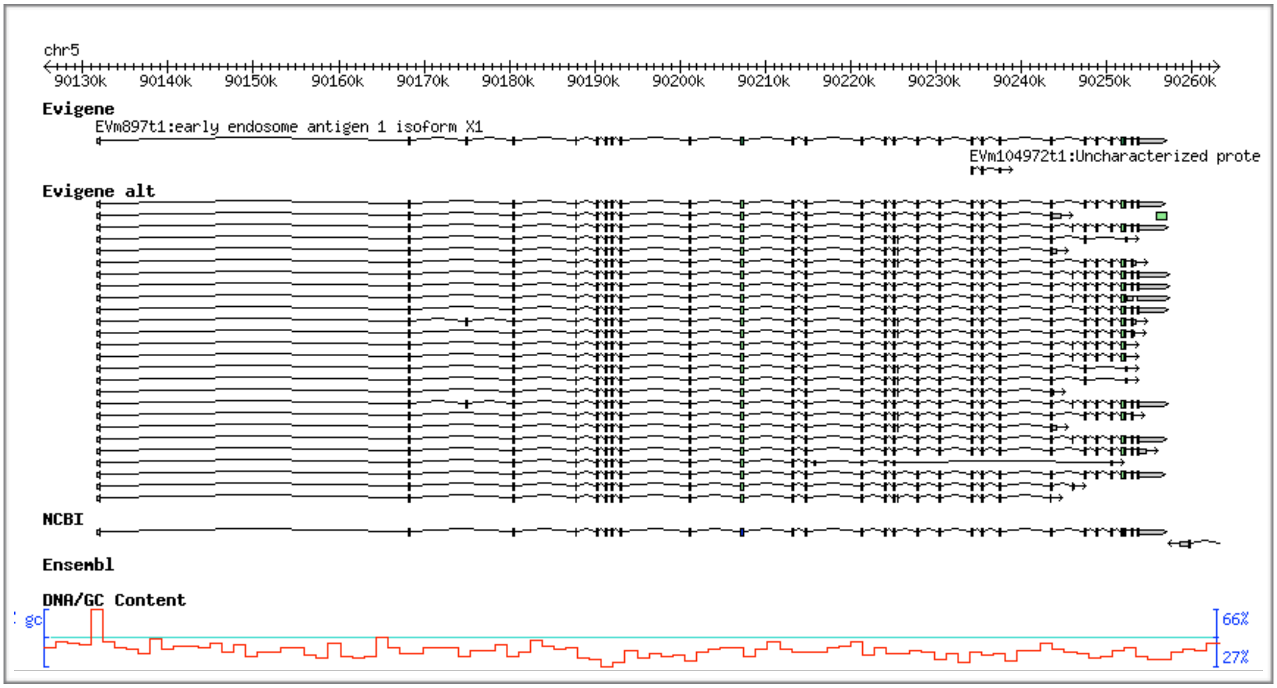

**Suppl. Figure 3, pigchr5gammatub**.   **Gene Name:** gamma-tubulin complex component 6;
**Notes:**  evg many alts, ncbi few alts, has all exons but all with truncated coding span (chr error likely), ensembl has most exons, but exon gaps at chr error span;  Models: human:NP_065194.2, 1819 aa, Pig Evigene: Susscr4EVm000660t1, 1831 align, NCBI:XP_020946990.1, 683 align,  Ensembl:ENSSSCP00000001040.3, 1203 align;

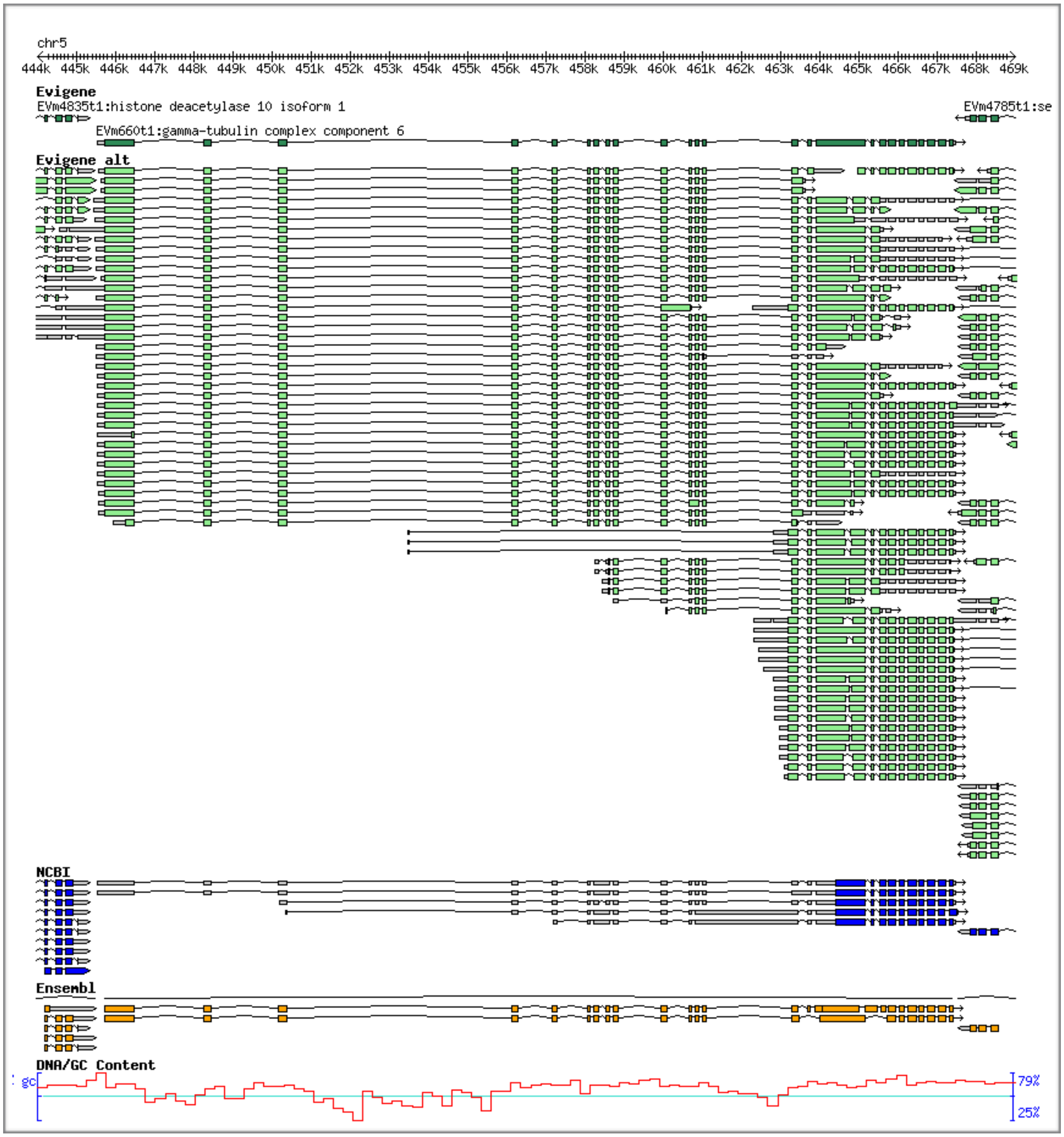

**Suppl. Figure 4, pigchr14trcoilcoil.** **Gene Name:** transforming acidic coiled-coil-containing protein 2; **Notes:** evg much longer model, many alts, more CDS exons, ncbi 1 valid shorter model, ensembl 1 valid even shorter model and 2 fragment models at 5' end of full gene; Models: human:XP_005269449.1, 2969 aa, Pig Evigene: Susscr4EVm000117t1, 2900 align, NCBI:NP_001245281.1, 1049 align, Ensembl:ENSSSCP00000058060.1, 1888 align

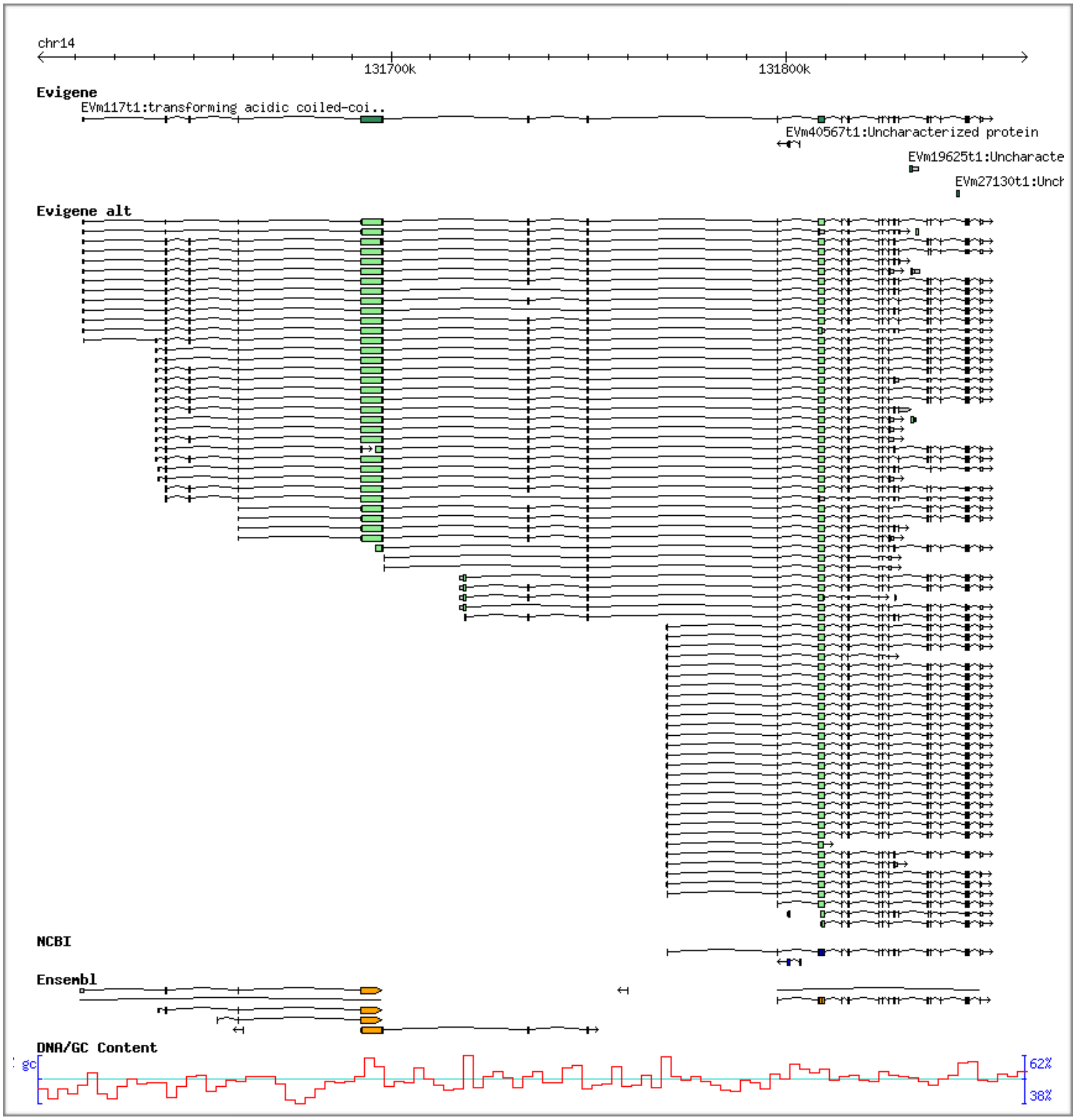

**Suppl. Figure 5, pigchr15titan.** **Gene Name:** titin, the longest protein at 36,000 aa, 108,000 cds bases in human, is a difficult gene to accurately model as 1/100,000 sequence errors for DNA or RNA will fragment this protein. **Notes:** evg and ncbi both have valid long model of 33,000 aa, ncbi is 3% longer; evg has many alts, ncbi 1 only; ensembl missed this largest gene entirely (has a few tiny fragments in span); Models: human:NP_001254479.2, 35991 aa, Pig Evigene: Susscr4EVm000001t1, 30600 align, NCBI:XP_020931560.1, 31391 align, Ensembl: missing;

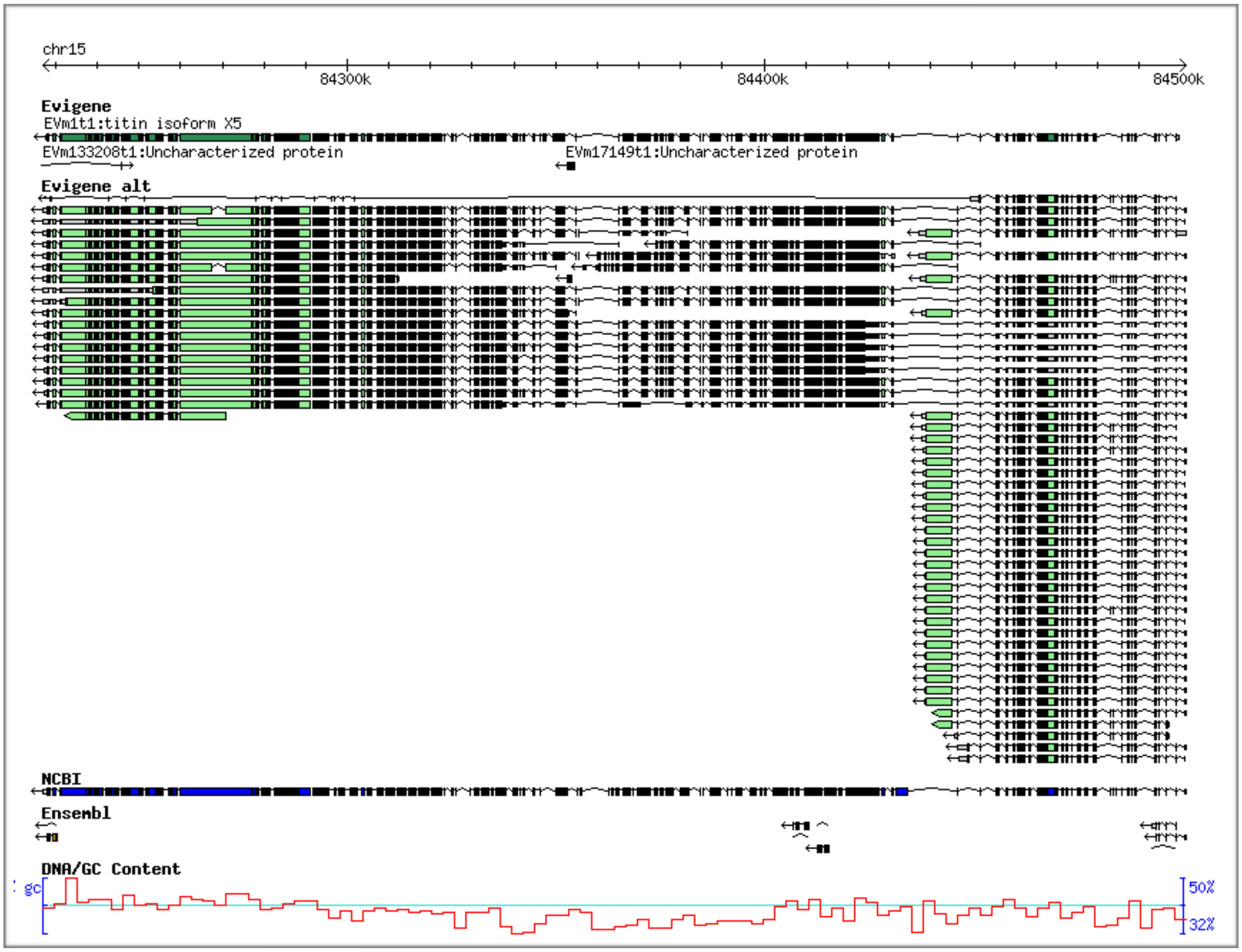

**Suppl. Figure 6, pigchr6medrnapol.** **Gene Name:** mediator of RNA polymerase II transcription subunit 25; **Notes:** evg full span and many alts, ncbi 1 model, missing 3' coding exon(s), ensembl 1 model + 1 fragment, missing coding exons; Models: human:XP_011525655.1, 796 aa, Pig Evigene: Susscr4EVm002758t2, 790 align, NCBI:XP_003127384.4, 715 align, Ensembl:ENSSSCP00000003469.3, 505 align

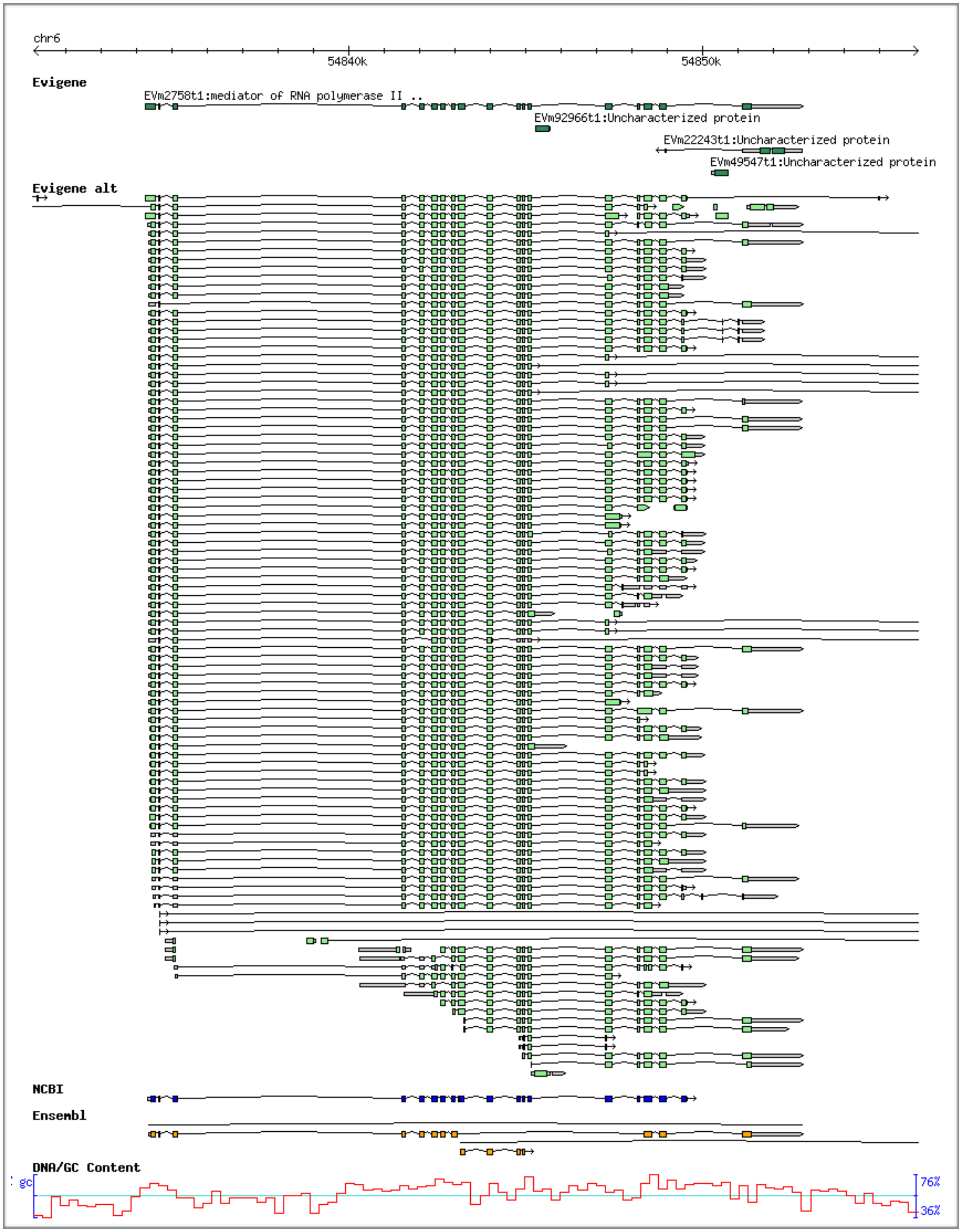

**Suppl. Figure 7, pigchr6microtube. Gene Name:** microtubule-actin cross-linking factor 1 ;
**Notes:** evg many alts is reason for >human align, ncbi 1 model valid but lacks some human exons, ensembl model is shorter fragment; Picture has cropped out 2/3 of Evigene alternates.
Models: human:XP_005270751.1, 7723 aa, Pig Evigene: Susscr4EVm000004t1, 7720 align, NCBI:XP_013833199.1, 6121 align, ENSSSCP00000038187.1, 6121 align

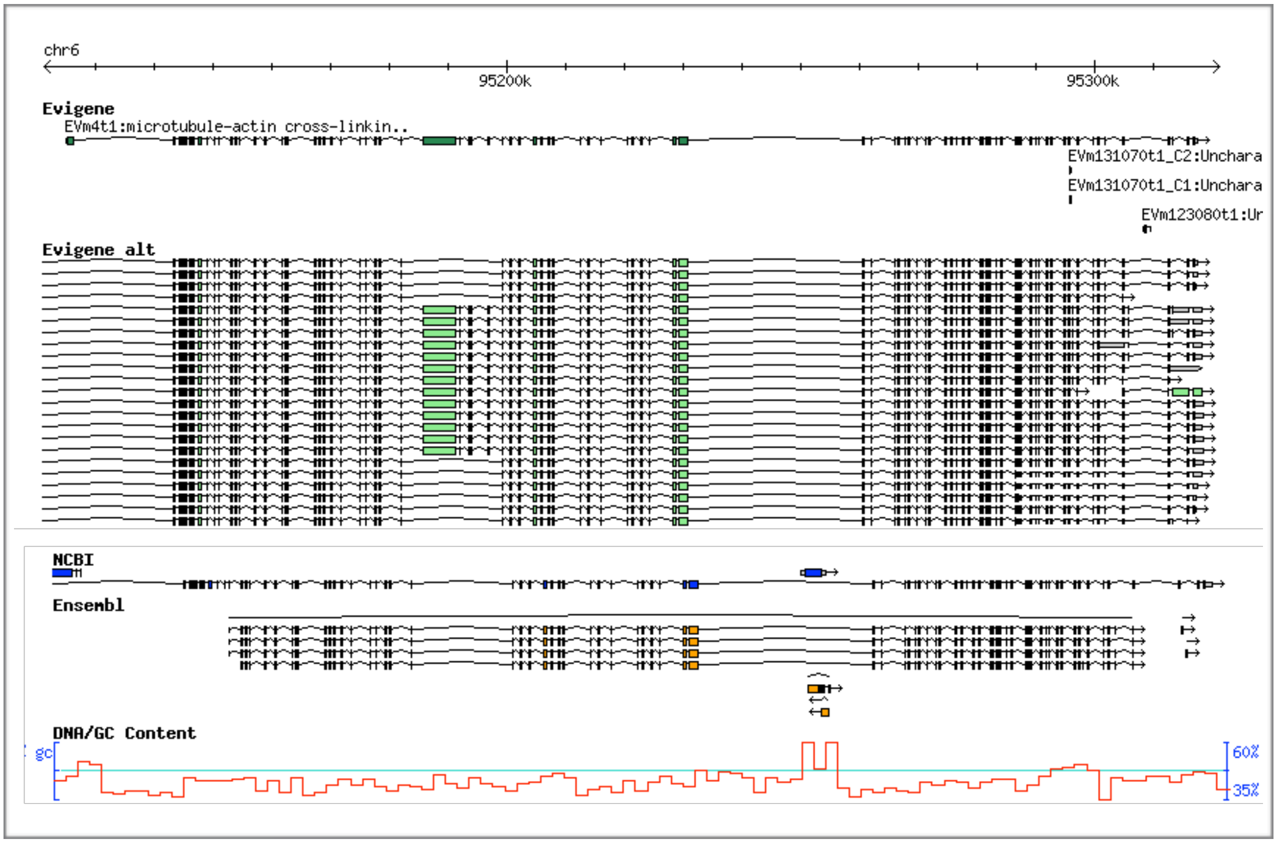

**Suppl. Figure 8, pigchr7unc79**. **Gene Name:** unc-79 homolog protein; **Notes:** evg and ncbi full models, ensembl 1 short model missing many CDS exons; Models human:XP_011535320.1, 2752, Pig Evigene:Susscr4EVm000169t1, 2699 align, NCBI:XP_020955291.1, 2734 align, Ensembl: missing

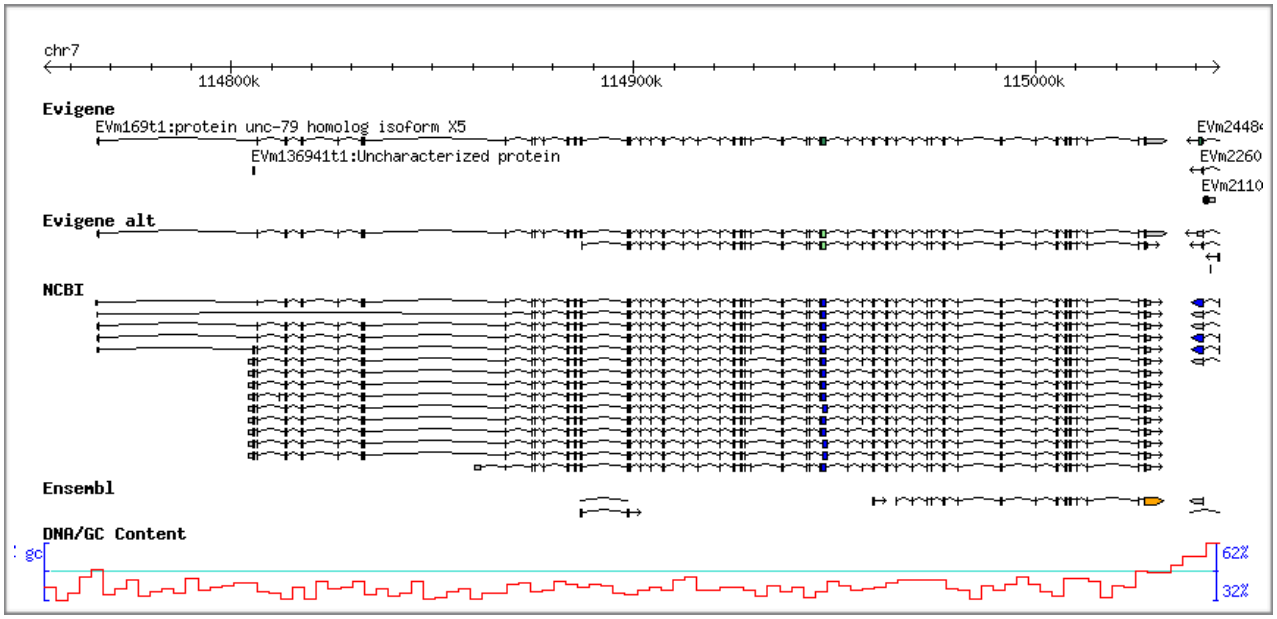

**Suppl. Figure 9, pigchr14nucrecptor**. **Gene Name:** nuclear receptor corepressor 2; **Notes:** evg many alts, more coding exons for greater human alignment, 1 model for ncbi and ensembl, ncbi has 2 more cds exons than ensembl; Picture has cropped out 2/3 of Evigene alternates. Models: human:NP_006303.4, 2514 aa, Pig Evigene: Susscr4EVm000084t1, 2504 align, NCBI:NP_001038041.1, 2464 align, Ensembl: ENSSSCP00000010425.2, 2465 align.

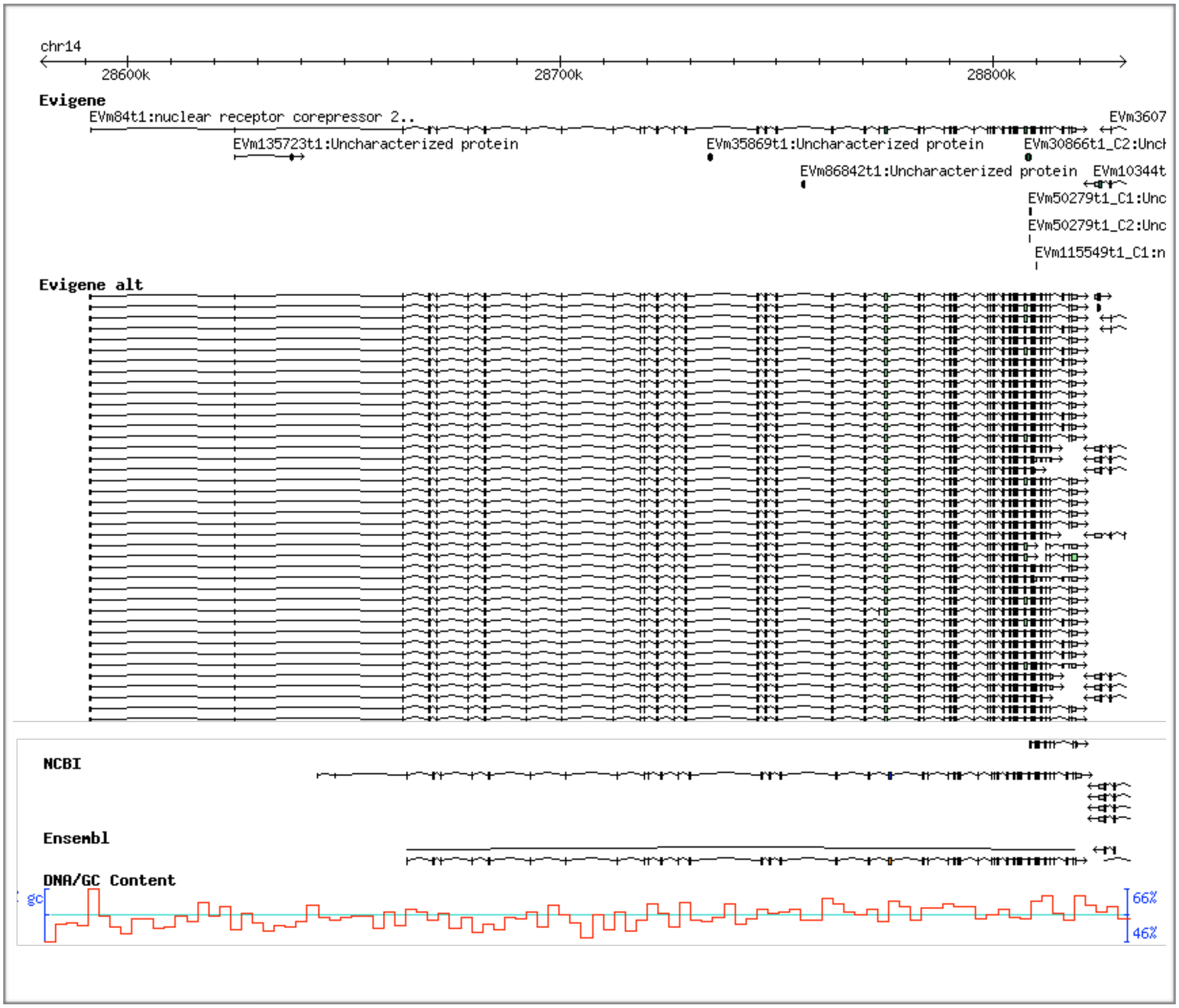

**Suppl. Figure 10, pigchr15strmuscle**. **Gene Name:** striated muscle preferentially expressed protein kinase; **Notes:** evg many alts, ncbi 1 model same as evg longest, ensembl fragment model missing many CDS exons; Models: human:XP_011508781.1, 3277 aa, Pig Evigene: Susscr4EVm000076t1, 3332 align, ncbi:XP_020930538.1, 3284 align, ENSSSCP00000036172.1, 844 align

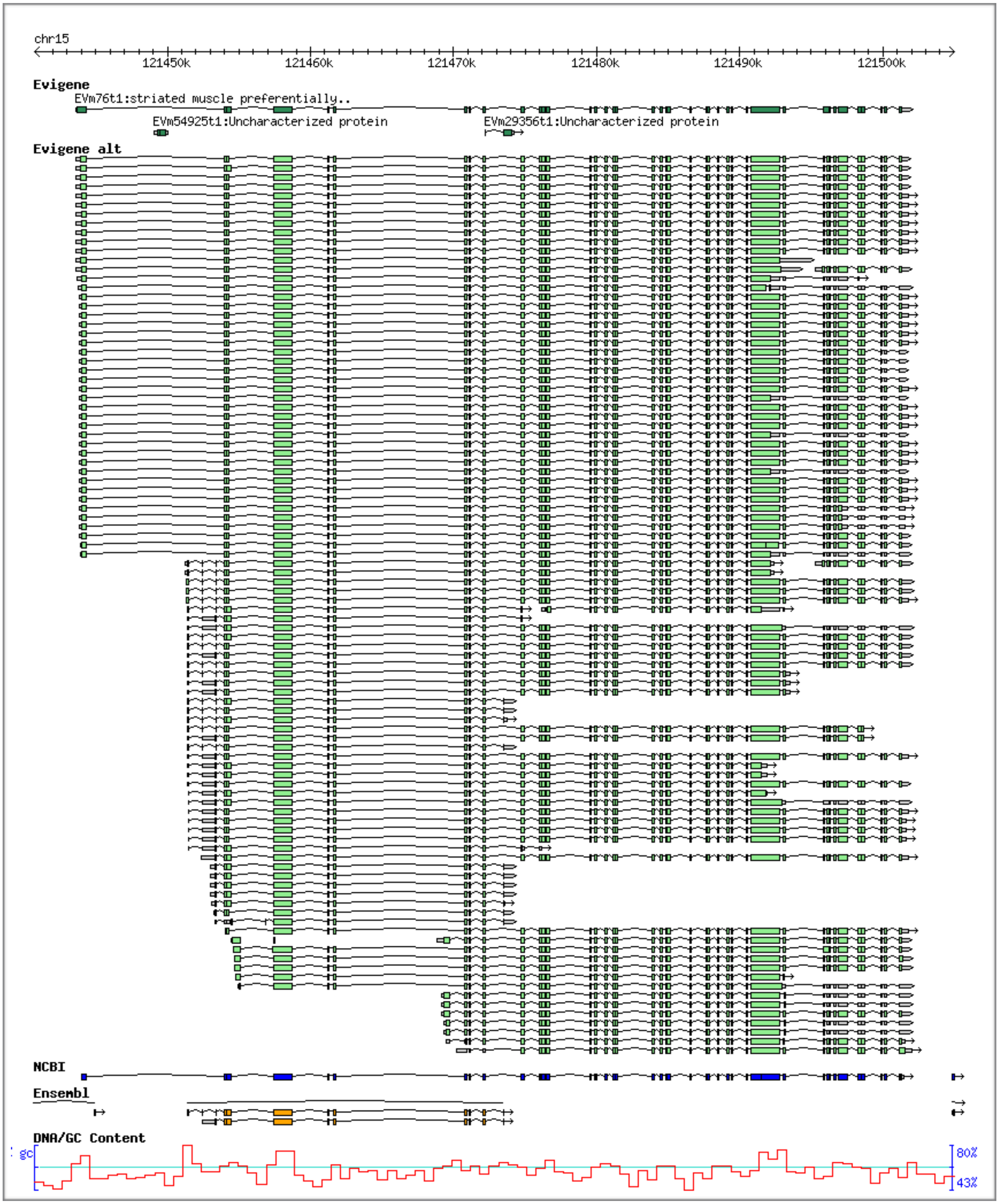
