## Supplemental Archive 4 for "Genes of the Pig, *Sus scrofa*, reconstructed with EvidentialGene": run_diginorm.pig1a.sh

```
#! /bin/bash
## env name=outname inpe=readpairs.fa datad=path/to/data this.sh
#PBS -N diginorm
#PBS -A PutAccountIdHere
#PBS -l nodes=1:ppn=16,walltime=39:55:00
#PBS -V

if [ "X" = "X$ncpu" ]; then ncpu=8; fi
if [ "X" = "X$maxmem" ]; then maxmem=32000; fi
if [ "X" = "X$datad" ]; then datad=/oasis/projects/nsf/ind114/ux455375/chrs/verts/sra2genes/evg17pig; fi
if [ "X" = "X$inspot" ]; then inspot='../spotfa/SRR6236879.fasta ../spotfa/SRR6236888.fasta ../spotfa/SRR6236889.fasta ../spotfa/SRR6236876.fasta ../spotfa/SRR6236881.fasta ../spotfa/SRR6236882.fasta'; fi
# options: what defaults?
if [ "X" = "X$kmer" ]; then kmer=25; fi
if [ "X" = "X$keep" ]; then keep=20; fi

dv="1a"; 

#** requires python2.6+ .. need this?
module add python

evigenes=/home/ux455375/bio/evigene/scripts

## /home/ux455375/bio/khmer/scripts == $HOME/bio/khmer ? or $HOME/bio/khmer/scripts
export PATH=/home/ux455375/bio/khmer/scripts:$PATH
export PYTHONPATH=/home/ux455375/bio/khmer/scripts/../python
normapp=/home/ux455375/bio/khmer/scripts/normalize-by-median.py

#  ncpu=2 # this is 1-cpu app, I think, may have changed.
kopt=""
outdir=spotfa
#old# outdir=rnasets

# FIXME: calc xhash from maxmem (in MB), for data size, along with $kmer and $keep
# 8e9 x 4 = 32Gb ; 12e9 x 4 = 48Gb; xhash=  1000000[mb2b]/4 * $maxmem
# xhash=8e9
xhash=$(( 250000 * $maxmem )) 

# output to rnasets/ folder of assembler inputs? or need tmp folder?
cd $datad/
mkdir $outdir
cd $outdir

echo "#START `date` " 

## inspot can be list..
echo $normapp $kopt -C $keep -x $xhash  -k $kmer  $inspot
$normapp $kopt -C $keep -x $xhash  -k $kmer  $inspot

echo "#DONE : `date`"
```
