## Supplemental Archive 4 for "Genes of the Pig, *Sus scrofa*, reconstructed with EvidentialGene": run_evgblastp.pig4321b.sh

```
#! /bin/bash
### env aaset=my.aa refaa=ref.aa datad=path/to/data qsub -q normal evgblastp.sh
#PBS -N evgblast
#PBS -A PutAccountIdHere
#PBS -l nodes=1:ppn=16,walltime=39:55:00
#PBS -V

if [ "X" = "X$ncpu" ]; then ncpu=10; fi
if [ "X" = "X$maxmem" ]; then maxmem=32000; fi
if [ "X" = "X$datad" ]; then datad=/oasis/projects/nsf/ind114/ux455375/chrs/verts/sra2genes/evg17pig4321mb; fi
if [ "X" = "X$aaset" ]; then aaset=pig4321b_okall.aa; fi
if [ "X" = "X$refaa" ]; then refaa=refset/refgenes18nc.aa; fi

evigenes=/home/ux455375/bio/evigene/scripts
# blastp/n/..:
export PATH=/home/ux455375/bio/ncbi/bin:$PATH

cd $datad/

qname=`basename $aaset .aa`
refnam=`basename $refaa .aa`

blopt="-evalue 1e-5"
odir=blout1$qname
mkdir $odir

echo "#START `date` " 

if [ ! -f $refaa.psq ]; then
  echo makeblastdb -dbtype prot -in $refaa -logfile $refaa.mblog
  makeblastdb -dbtype prot -in $refaa -logfile $refaa.mblog
fi

if [ ! -f $aaset.split.1.fa ]; then
 pindir=`dirname $aaset`
 splitsize=`grep -v '^>' $aaset | wc -c | sed 's/ .*//' `
 splitbp=$(( $splitsize / $ncpu ))
 $evigenes/splitMfasta.pl --outputpath=$pindir --nparts $ncpu --minsize=$splitbp $aaset
fi

qset=`/bin/ls $aaset.split.*.fa`

for qfile in $qset
{
  qnamspl=`basename $qfile .fa`
  onam=$odir/$refnam-$qnamspl
  echo blastp $blopt -outfmt 7 -db $refaa -query $qfile -out $onam.blastp
  blastp $blopt -outfmt 7 -db $refaa -query $qfile -out $onam.blastp  &
}

wait

rqname=$refnam-$qname
aablast=$rqname.blastp
aabltab=$rqname.btall

cat $odir/$rqname.*.blastp > $aablast
/bin/rm $qset
echo $odir is temp blast output, check for completeness then erase
echo /bin/rm -r $odir

env oid=1 off=1 $evigenes/prot/aaqual.sh $aaset
env oid=1 off=1 $evigenes/prot/aaqual.sh $refaa
mbaopts="-tall -aasize $aaset.qual,$refaa.qual"

$evigenes/makeblastscore3.pl $mbaopts $aablast > $aabltab 

# ADD here STEP9. namegenes from ref names, for annotation .. but need ref.names collection 
echo $evigenes/prot/namegenes.pl -blast $aabltab -refnames $refaa.names -out $qname.names
if [ -f $refaa.names ]; then
  $evigenes/prot/namegenes.pl -blast $aabltab -refnames $refaa.names -out $qname.names
else 
  echo MISSING $refaa.names for  namegenes.pl  .. -out $qname.names
fi

gzip --fast $aablast

echo "#DONE : `date`"
```
