## Supplemental Archive 4 for "Genes of the Pig, *Sus scrofa*, reconstructed with EvidentialGene": run_evgpubset.pig4321b.sh

```
#! /bin/bash
### env idprefix=MysppEGm trclass=myspp.trclass datad=`pwd` qsub -q shared run_evgmrna2tsa.sh
#PBS -N evgpubset
#PBS -A PutAccountIdHere
#PBS -l nodes=1:ppn=16,walltime=39:55:00
#PBS -V

if [ "X" = "X$ncpu" ]; then ncpu=10; fi
if [ "X" = "X$maxmem" ]; then maxmem=32000; fi
if [ "X" = "X$datad" ]; then datad=/oasis/projects/nsf/ind114/ux455375/chrs/verts/sra2genes/evg17pig4321mb; fi
if [ "X" = "X$trclass" ]; then trclass=pig4321b.trclass; fi
#FIXME:if [ "X" = "X$species" ]; then species=Sus_scrofa;Sus_scrofa_domesticus; fi
if [ "X" = "X$species" ]; then species=Sus_scrofa; fi
if [ "X" = "X$idprefix" ]; then idprefix=SusscrEVm; fi

export evigenes=/home/ux455375/bio/evigene/scripts
export PATH=/home/ux455375/bio/ncbi/bin:$PATH

opts="-debug  -dropshow -skipdropseq -NCPU $ncpu"

## .. these are now read via sra_result.csv, species => idprefix
if [ "X" != "X$idprefix" ]; then opts="$opts -idprefix $idprefix"; fi
#NO#if [ "X" = "X$vectrim" ]; then opts="$opts -novectrim"; fi
if [ "X" != "X$tbl2asn" ]; then opts="$opts -runtbl2asn"; fi
if [ "X" != "X$species" ]; then spp=`echo $species | sed 's/ /_/g;'`; opts="$opts -species=$spp"; fi
if [ "X" = "X$names" ]; then names=`echo $trclass | sed 's/.gz//; s/\.trclass/.names/;'`; fi
if [ -s $names ]; then opts="$opts -names $names"; fi

cd $datad/

echo "#START `date` " 
echo $evigenes/evgmrna2tsa2.pl  $opts -log -class $trclass
if [ ! -f $trclass ]; then echo "ERR: missing -class $trclass"; exit -1; fi
$evigenes/evgmrna2tsa2.pl  $opts -log -class $trclass
echo "#DONE : `date`"
```
