## Supplemental Archive 4 for "Genes of the Pig, *Sus scrofa*, reconstructed with EvidentialGene": run_evgsra2genes.sh

```
#! /bin/bash
### env sratable=sraset.csv datad=`pwd` ncpu=16 qsub -q normal run_evgsra2genes.sh
#PBS -N evgsra2genes
#PBS -A PutAccountIdHere
#PBS -l nodes=1:ppn=16,walltime=39:55:00
#PBS -V

#. APPS_PATH=XSEDE
#. APPS_PATH=MODULE
APPS_PATH=MAC

if [ "X" = "X$ncpu" ]; then ncpu=8; fi
if [ "X" = "X$maxmem" ]; then maxmem=64000; fi
if [ "X" = "X$datad" ]; then echo "ERROR: missing datad=/path/to/data"; exit -1; fi
if [ "X" = "X$sratable" ]; then echo "ERROR: missing sratable=/path/to/data"; exit -1; fi

# dgg home
if [ $APPS_PATH = "MAC" ]; then
  bioapps=/bio/apps
  evigenes=$bioapps/evigene/scripts
  srabin=$bioapps/sratools/sratoolkit.2.8.1-2-mac64/bin
  velobin=$bioapps/rnaseq/velvs/velo120/velbin1  # fixme multi kmer binaries
  idbabin=$bioapps/rnaseq/idba/idba-1.1.1/bin
  soapbin=$bioapps/rnaseq/soaptr/SOAPdenovo-Trans-r104/bin  # fixme binaries
  trinbin=$bioapps/rnaseq/trin12/trinityrnaseq_r2012-10-05/ # fixme binaries
  xnrbin=$bioapps/exonerate/bin
  cdhitbin=$bioapps/cdhit17/bin
  ncbibin=$bioapps/ncbix/bin
  kmerbin=$bioapps/khmer/scripts
  gmapbin=$bioapps/gmap1701/bin
fi

if [ $APPS_PATH = "MODULE" ]; then
  # dont know if anyone has all these in module installations..
  module add blast sratoolkit
  module add evigene
  module add exonerate cdhit khmer
  module add gmap_gsnap
  module add oases soaptrans idba trinity
fi

# XSEDE .sdsc.edu
if [ $APPS_PATH = "XSEDE" ]; then
  bioapps=$HOME/bio
  evigenes=$bioapps/evigene/scripts
  # NOTE need current sratoolkit281 for web fetch by SRR id
  srabin=$bioapps/sratoolkit/sratoolkit281/bin
  #  velvet: fixme multi kmer binaries, bin4 = 151mer; bin2 = 99mer
  velobin=$bioapps/velvet1210/bin  
  idbabin=$bioapps/idba/bin
  soapbin=$bioapps/soaptrans103 
  trinbin=$bioapps/trinity
  xnrbin=$bioapps/exonerate/bin
  cdhitbin=$bioapps/cdhit466/bin   
  ncbibin=$bioapps/ncbi/bin
  kmerbin=$bioapps/khmer/scripts
  gmapbin=$bioapps/gmap/bin
fi

addpath=$srabin:$ncbibin:$velobin:$idbabin:$soapbin:$trinbin
addpath=$evigenes:$xnrbin:$cdhitbin:$gmapbin:$kmerbin:$addpath
export PATH=$addpath:$PATH

evopts="-NCPU $ncpu -MAXMEM $maxmem -log -debug"
## yes, add opt -runstep start7 start at evgreduce step7, with merge/trsets/ 
if [ "X" != "X$runsteps" ]; then evopts="$evopts -runsteps $runsteps"; fi
if [ "X" != "X$name" ]; then evopts="$evopts -runname $name"; fi

cd $datad
echo $evigenes/evgpipe_sra2genes.pl $evopts -SRAtable $sratable
$evigenes/evgpipe_sra2genes.pl $evopts -SRAtable $sratable

##--------------------------------------------
## PATHS .. see evgpipe_sra2genes:findapp() for needed paths
#   findapp('fastq-dump');  
#   findapp('idba_tran', 1);
#   findapp('velveth', 1); # revise velbin compile version names: vel{h,g},oases_NNNmer for max kmer
#   findapp('SOAPdenovo-Trans-127mer', 1);
#   findapp('Trinity', 1);
#   findapp('fastanrdb', 1); 
#   findapp('cd-hit-est', 1); 
#   findapp('blastn', 1); # also vecscreen tbl2asn maybe
#   findapp('gmap', 1)
#
```
