## Supplemental Archive 4 for "Genes of the Pig, *Sus scrofa*, reconstructed with EvidentialGene": run_evgsra2genes_scripts.txt

# run_evgsra2genes, script of scripts
#-------------------
# These are command-line calls typed by the author to execute SRA2Genes for the Pig Gene Assembly,
# on a cluster system, comet.sdsc.edu of XSEDE, in serial fashion.  
# First run_evgsra2genes.sh is input of SRA ID table of a project RNA run entries, and results in
#  SRA data fetching, and new scripts to run RNA assemblers on cluster.
# Subsequent invocations of run_evgsra2genes.sh, and $evigene/scripts/evgpipe_sra2genes.pl,
#  then check current data results, and write new cluster scripts for the next data processing steps.
# srun_comet.sh is the cluster system sbatch script, which runs the prog=... genome script,
# in data directory datad=..., with given environment parameters (name=, sratable=, ..)
#-------------------

# sample data tables from NCBI SRA queries by project: 
pig1a=srapigPRJNA416432.csv
pig2b=srapigPRJNA353772.csv	
pig3c=srapigPRJEB8784.csv	
pig4e=srapigPRJNA255281.csv	

# sample pig1a, first run of SRA2Genes
env name=pig1a sratable=srapigPRJNA416432.csv datad=`pwd` prog=./run_evgsra2genes.sh sbatch srun_shared.sh

# sample pig3c, first run of SRA2Genes
env name=pig3c sratable=srapigPRJEB8784.csv datad=`pwd` prog=./run_evgsra2genes.sh sbatch srun_shared.sh

# scripts written by SRA2Genes for SRA data fetched and assemblers; send to cluster batch with prog=scriptname
env prog=./runidba.pig1a.sh datad=`pwd` sbatch srun_comet.sh
env prog=./runsoap.pig1a.sh datad=`pwd` sbatch srun_comet.sh
env prog=./runvelo.pig1a.sh datad=`pwd` sbatch srun_comet.sh
env prog=./runtrin.pig1a.sh datad=`pwd` sbatch srun_comet.sh
env prog=./run_diginorm.pig1a.sh datad=`pwd` sbatch srun_share8m48h40.sh

# repeat run_evgsra2genes with same options, after first step, for next cluster batch scripts
env name=pig1a sratable=srapigPRJNA416432.csv datad=`pwd` prog=./run_evgsra2genes.sh sbatch srun_shared.sh

# following assembler runs, SRA2Genes collects outputs for input to next step, transcripts reduction
env prog=./run_tr2aacds.pig1a.sh datad=`pwd` sbatch srun_share8m48h40.sh

# after assembly+reduction of four SRA projects (pig1,2,3,4), merge gene sets start at Step7 of SRA2Genes
# .. pig4321ma, catenate tr2aacds okayset results as trset inputs to merged set
# .. use new start point of run_evgsra2genes
env name=pig4321b runsteps=start7 datad=`pwd` prog=../run_evgsra2genes.sh sbatch ../srun_shared.sh

# annotate, vector/contam checks, public set results with further steps of SRA2Genes
env prog=./run_tr2aacds.pig4321b.sh datad=`pwd` sbatch srun_share8m48h40.sh
env prog=./run_evgblastp.pig4321b.sh  datad=`pwd` sbatch srun_comet.sh
env prog=./run_evgtrimvec.pig4321b.sh datad=`pwd` sbatch srun_comet.sh

env idprefix=Susscr1EVm prog=./run_evgpubset.pig4321b.sh datad=`pwd` sbatch srun_share8m48h40.sh 
