## Supplemental Archive 4 for "Genes of the Pig, *Sus scrofa*, reconstructed with EvidentialGene": run_evgtrimvec.pig4321b.sh

```
#! /bin/bash
### env trclass=myspp.trclass datad=`pwd` qsub -q shared run_evgtrimvec.sh
#PBS -N evgtrimvec
#PBS -A PutAccountIdHere
#PBS -l nodes=1:ppn=16,walltime=39:55:00
#PBS -V

if [ "X" = "X$ncpu" ]; then ncpu=10; fi
if [ "X" = "X$maxmem" ]; then maxmem=32000; fi
if [ "X" = "X$datad" ]; then datad=/oasis/projects/nsf/ind114/ux455375/chrs/verts/sra2genes/evg17pig4321mb; fi
if [ "X" = "X$trclass" ]; then trclass=pig4321b.trclass; fi
#bad#if [ "X" = "X$mrna" ]; then mrna=P_MRNA; fi
## MINSIZE is an option, transcripts smaller are DROPped, 200 is common but some valid ortho are smaller ..
if [ "X" = "X$MINSIZE" ]; then MINSIZE=180; fi

export evigenes=/home/ux455375/bio/evigene/scripts
export vecscreen=/home/ux455375/bio/ncbi/bin/vecscreen
export PATH=/home/ux455375/bio/ncbi/bin:$PATH

#b# trimopts="-nodeferupdate -MINSIZE $MINSIZE -NCPU $ncpu -log"
#b# namepath=`echo $mrna | sed 's/\.mrna//; s/\.fasta//; s/\.fa//; s/\.tr//;'`
trimopts="-deferupdate -MINSIZE $MINSIZE -NCPU $ncpu -log"
namepath=`echo $trclass | sed 's/\.trclass//;'`

cd $datad/
echo "#START `date` " 

if [ -f $namepath.trimvec_done ]; then
  echo "# $namepath.trimvec_done"
else
  echo $evigenes/rnaseq/asmrna_trimvec.pl $trimopts -trclass $trclass
  $evigenes/rnaseq/asmrna_trimvec.pl $trimopts -trclass $trclass
fi

echo "#DONE : `date`"
```
