## Supplemental Archive 4 for "Genes of the Pig, *Sus scrofa*, reconstructed with EvidentialGene": run_gmapgenes.pig4321b.sh

```
#! /bin/bash
### env  mrna=publicset/prname.mrna_pub.fa datad=`pwd` qsub -q normal run_gmapgenes.sh
#PBS -N gmapgenes
#PBS -A PutAccountIdHere
#PBS -l nodes=1:ppn=16,walltime=39:55:00
#PBS -V

if [ "X" = "X$ncpu" ]; then ncpu=10; fi
if [ "X" = "X$maxmem" ]; then maxmem=32000; fi
if [ "X" = "X$datad" ]; then datad=/oasis/projects/nsf/ind114/ux455375/chrs/verts/sra2genes/evg17pig4321mb; fi
if [ "X" = "X$mrna" ]; then mrna=publicset/pig4321b.mrna_pub.fa; fi
if [ "X" = "X$genome" ]; then genome=genome/chrpig11c.fa.gz; fi
if [ "X" = "X$name" ]; then name=`basename $mrna .mrna | sed 's/.gz//; s/.mrna_pub.fa//; s/\.fa.*$//'`; fi

export evigenes=/home/ux455375/bio/evigene/scripts
export PATH=P_NCBIBIN:$PATH
export PATH=/opt/biotools/gmap_gsnap/bin:$PATH
#-----

gmapopt="--suboptimal-score=1 --min-intronlength=18 --microexon-spliceprob=1.0 -S"
gbuildopt=""
# gmap doesnt use suffix array, gsnap does, aids gsnap speed, slows gmap_build, not required
# gbuildopt="--build-sarray=0 --no-sarray"

gdb=genome/gmap
gnogz=`echo $genome | sed 's/.gz//;'`
isgz=1; if [ $gnogz = $genome ]; then isgz=0; fi
gename=`basename $genome .fa.gz | sed 's/.gz//; s/\.fa.*//;'`

cd $datad/

echo "#START `date` " 
if [ ! -s $mrna ]; then echo "missing mrna = $mrna"; exit -1; fi

if [ ! -d $gdb/$gename ]; then
  if [ ! -s $genome ]; then echo "missing genome.fasta = $genome"; exit -1; fi
  if [ ! -d $gdb ]; then mkdir -p $gdb; fi
  if [ $isgz = 1 ]; then  gbuildopt="$gbuildopt --gunzip"; fi
  echo gmap_build $gbuildopt -d $gename -D $gdb $genome
  gmap_build $gbuildopt -d $gename -D $gdb $genome
fi

namege=$name-$gename
nameout=$namege.gmap.out

gnogz=`echo $mrna | sed 's/.gz//;'`
isgz=1; if [ $gnogz = $mrna ]; then isgz=0; fi
echo gmap $gmapopt -d $gename -D $gdb --nthreads=$ncpu $mrna TO $nameout
if [ $isgz = 1 ]; then
  gunzip -c $mrna | gmap $gmapopt -d $gename -D $gdb --nthreads=$ncpu > $nameout
else 
  gmap $gmapopt -d $gename -D $gdb --nthreads=$ncpu $mrna > $nameout
fi

eggopt="-nocututrchim -nopath -noerrspan -best=0 -intron=-1"
annopt="";  
gann=`echo $mrna | sed 's/\.mrna.*/.ann.txt/;'` 
if [ ! -f $gann ]; then
  gann=`echo $mrna | sed 's/\.mrna.*/.names/;'` 
fi
if [ -f $gann ]; then annopt="-names $gann"; fi

if [ -s $nameout ]; then
  echo cat $nameout P gmap2evgff.pl P evganngff.pl $annopt TO  $namege.evgmap.gff
  cat $nameout | $evigenes/genes/gmap2evgff.pl $eggopt -src evgmap \
   | $evigenes/genes/evganngff.pl $annopt -genes stdin > $namege.evgmap.gff
  $evigenes/ests/gff2aligntab.pl < $namege.evgmap.gff | sort -k1,1 -k2,2n -k4,4 > $namege.align.tab
  gzip --fast $nameout
fi

echo "#DONE : `date`"
```
