## Supplemental Archive 4 for "Genes of the Pig, *Sus scrofa*, reconstructed with EvidentialGene": run_tr2aacds.pig1a.sh

```
#! /bin/bash
### env datad=path/to/data trset=myspecies_all.tr.gz qsub -q normal runtr2cds.sh
#PBS -N tr2aacds
#PBS -A PutAccountIdHere
#PBS -l nodes=1:ppn=16,walltime=39:55:00
#PBS -V

if [ "X" = "X$ncpu" ]; then ncpu=10; fi
if [ "X" = "X$maxmem" ]; then maxmem=32000; fi
if [ "X" = "X$datad" ]; then datad=/oasis/projects/nsf/ind114/ux455375/chrs/verts/sra2genes/evg17pig; fi
if [ "X" = "X$trset" ]; then trset='pig1a.tr'; fi
if [ "X" = "X$name" ]; then name=`basename $trset .tr | sed 's/\.fa.*$//'`; fi

evigenes=/home/ux455375/bio/evigene/scripts
evapp=$evigenes/prot/tr2aacds2.pl
traopts="-tidy -log -debug"
if [ "X" != "X$opt" ]; then traopts="$traopts $opt"; fi
#^ test new opt=-small/-nosmall for 2level reduction, aablast between

# cd-hit-est/aa
# export PATH=$HOME/bio/cdhit/bin:$PATH
export PATH=/home/ux455375/bio/cdhit466/bin:$PATH
# export fastanrdb=$HOME/bio/exonerate/bin/fastanrdb
export fastanrdb=/home/ux455375/bio/exonerate/bin/fastanrdb
# blastn:
export PATH=/home/ux455375/bio/ncbi/bin:$PATH

cd $datad/

echo "#START `date` " 
echo $evapp -NCPU $ncpu -MAXMEM $maxmem $traopts -cdna $trset
$evapp -NCPU $ncpu -MAXMEM $maxmem $traopts -cdna $trset
echo "#DONE : `date`"
```
