## Supplemental Archive 4 for "Genes of the Pig, *Sus scrofa*, reconstructed with EvidentialGene": runidba.pig1a.sh

```
#! /bin/bash
## env name=outname inpe=readpairs.fa datad=path/to/data this.sh
#PBS -N idba_trans
#PBS -A PutAccountIdHere
#PBS -l nodes=1:ppn=16,walltime=39:55:00
#PBS -V

if [ "X" = "X$ncpu" ]; then ncpu=8; fi
if [ "X" = "X$maxmem" ]; then maxmem=32000; fi
if [ "X" = "X$datad" ]; then datad=/oasis/projects/nsf/ind114/ux455375/chrs/verts/sra2genes/evg17pig; fi
if [ "X" = "X$inpe" ]; then inpe='rnasets/sBn1l1SRR6236889.fa2'; fi
if [ "X" = "X$name" ]; then name=sBn1l1SRR6236889; fi

dv="1a"; # versnum, add opt
traopts="--mink 27 --maxk 117 --step 10 --max_isoforms 9";

evigenes=/home/ux455375/bio/evigene/scripts
idba_tran=/home/ux455375/bio/idba/bin/../bin2/idba_tran
export PATH=/home/ux455375/bio/idba/bin/../bin2:$PATH

#o# outdir=tridba$name$dv
outdir=tridba${dv}_$name
cd $datad/

echo "#START  `date`"
echo "$idba_tran $traopts --read $inpe --num_threads $ncpu --out $outdir"
$idba_tran $traopts --read $inpe --num_threads $ncpu --out $outdir

## add cleanups, where contig.fa  is ok .. all graph files?
## kmer [big]; align-* component-* connection-* contig-$ks.fa contig-info-*.fa graph-*.fa local-contig-*.fa transcript-path-*
## leave unzipped transcript-*.fa contig.fa 
if [ -f $outdir/contig.fa ]; then
  cd $outdir
  gzip --fast kmer &
  gzip --fast align-* &
  gzip --fast graph-* &
  gzip --fast contig-*.fa local-contig-*.fa transcript-path-* component-* connection-* &
  cd ../
fi

echo "#DONE : `date`"
```
