## Supplemental Archive 4 for "Genes of the Pig, *Sus scrofa*, reconstructed with EvidentialGene": runsoap.pig1a.sh

```
#! /bin/bash
## env name=outname inpe=readpairs.fa datad=path/to/data this.sh
#PBS -N soap_trans
#PBS -A PutAccountIdHere
#PBS -l nodes=1:ppn=16,walltime=39:55:00
#PBS -V

if [ "X" = "X$ncpu" ]; then ncpu=8; fi
if [ "X" = "X$maxmem" ]; then maxmem=32000; fi
if [ "X" = "X$datad" ]; then datad=/oasis/projects/nsf/ind114/ux455375/chrs/verts/sra2genes/evg17pig; fi
if [ "X" = "X$inpe" ]; then inpe='rnasets/sBn1l1SRR6236889_1.fa'; fi
if [ "X" = "X$name" ]; then name=sBn1l1SRR6236889; fi

dv="1a"; 
outdir=trsoap${dv}_$name
onamep=$name$dv

INSIZE=200 #or 200
INSIZESD=50 # or 200SD  
# F: gapfil; M: merge strength 1=def
sopt="-p $ncpu -F -t 99"

kset_example="31 27 25 35 45 55"
kset="25 31 37 43 49 55 61 67 73 "

evigenes=/home/ux455375/bio/evigene/scripts
export PATH=/home/ux455375/bio/soaptrans103:$PATH

cd $datad/
mkdir $outdir

# write soap.config .. inpe presumed local path rel to datad
echo "max_rd_len=150" > $outdir/rdconfig 
for fa1 in $inpe; do {
  fa2=`echo $fa1 | sed 's/_1/_2/;'`
  if [ $fa2 = $fa1 ]; then continue; fi
  # nogz=`echo $fa1 | sed 's/\.gz//;'` # skip this, require ungzipped data
  cat >> $outdir/rdconfig <<EOT

[LIB]
rank=1
asm_flag=3
avg_ins=$INSIZE
f1=$datad/$fa1
f2=$datad/$fa2

EOT

} done

cd $outdir
echo "#START `date` " 

for kmer in $kset; do {
  odir=sod$kmer
  outname=so${onamep}.k$kmer
  mkdir $odir
  if [ $kmer -lt 32 ]; then
    SOAPdenovo-Trans-31mer all -s rdconfig -o $outname -K $kmer $sopt
  else 
    SOAPdenovo-Trans-127mer all -s rdconfig -o $outname -K $kmer $sopt
  fi

  mv $outname* $odir/
  if test -f $odir/$outname.contig ; then
    rm $odir/$outname.{readOnContig,readInGap,ctg2Read,preArc,vertex,edge.gz,Arc,updated.edge}
    gzip --fast $odir/$outname.{scafSeq,contig}
  fi

} done

echo "#DONE : `date`"
```
