## Supplemental Archive 4 for "Genes of the Pig, *Sus scrofa*, reconstructed with EvidentialGene": runtrin.pig1a.sh

```
#! /bin/bash
## env name=outname inpe=readpairs.fa datad=path/to/data this.sh
#PBS -N trinity
#PBS -A PutAccountIdHere
#PBS -l nodes=1:ppn=16,walltime=39:55:00
#PBS -V

if [ "X" = "X$ncpu" ]; then ncpu=8; fi
if [ "X" = "X$maxmem" ]; then maxmem=32000; fi
if [ "X" = "X$datad" ]; then datad=/oasis/projects/nsf/ind114/ux455375/chrs/verts/sra2genes/evg17pig; fi
if [ "X" = "X$inpe" ]; then inpe='../rnasets/sBn1l1SRR6236889_1.fa'; fi
if [ "X" = "X$name" ]; then name=`basename $inpe _1.fa | sed 's/\.fa.*$//'`; fi

dv="1a"; 

## what extras are  needed?
module add bowtie2 samtools

evigenes=/home/ux455375/bio/evigene/scripts
export PATH=/home/ux455375/bio/trinity:$PATH

infa1=$inpe
infa2=`echo $inpe | sed 's/_1/_2/;'`
outdir=trtrin${dv}_$name

# --output $outdir .. must include 'trinity' in the name .. dont use
# trin.maxmem in Gb of RAM WITH 'G' appended; evigene maxmem == Mb
tmaxmem=$(( $maxmem / 1000 )); tmaxmem="${tmaxmem}G";
topts="--CPU $ncpu --max_memory $tmaxmem --no_normalize_reads --SS_lib_type RF"

cd $datad/
mkdir $outdir
cd $outdir

echo "#START `date` " 
echo /home/ux455375/bio/trinity/Trinity $topts --seqType fa --left $infa1 --right $infa2
/home/ux455375/bio/trinity/Trinity $topts --seqType fa --left $infa1 --right $infa2
echo "#DONE : `date`"
```
