## Supplemental Archive 4 for "Genes of the Pig, *Sus scrofa*, reconstructed with EvidentialGene": runvelo.pig1a.sh

```
#! /bin/bash
## env name=outname inpe=readpairs.fa datad=path/to/data this.sh
#PBS -N velvetoases
#PBS -A PutAccountIdHere
#PBS -l nodes=1:ppn=16,walltime=39:55:00
#PBS -V

# more than 8 cpu bad, overuses memory ..
if [ "X" = "X$ncpu" ]; then ncpu=8; fi
if [ "X" = "X$maxmem" ]; then maxmem=32000; fi
if [ "X" = "X$datad" ]; then datad=/oasis/projects/nsf/ind114/ux455375/chrs/verts/sra2genes/evg17pig; fi
if [ "X" = "X$inpe" ]; then inpe='../rnasets/sBn1l1SRR6236889_1.fa ../rnasets/sBn1l1SRR6236889_2.fa'; fi
if [ "X" = "X$name" ]; then name=sBn1l1SRR6236889; fi

dv="1a"; 
outdir=trvelo${dv}_$name
INSIZE=200 #or 200
INSIZESD=50 # or 200SD  

# kset_example="95 85 75 71 65 63 55 45 35"
#orig:kset="145 135 125 115 105 95 85 75 79 65 55 45 35 "
kset="125 115 105 95 85 75 79 65 55 45 35 "

evigenes=/home/ux455375/bio/evigene/scripts
velbin=/home/ux455375/bio/velvet1210/bin2
## FIXME appbin variants... use /home/ux455375/bio/velvet1210/bin2/../bin2 /home/ux455375/bio/velvet1210/bin2/../bin4 P_A4BIN ?
#99<k<=155 ? bin4/
velbin4=/home/ux455375/bio/velvet1210/bin2/../bin4
#61<k<=99# 
velbin2=/home/ux455375/bio/velvet1210/bin2/../bin2
#k<=61# 
velbin1=/home/ux455375/bio/velvet1210/bin2
##......................

export OMP_NUM_THREADS=$ncpu

vopth=""
iopts="-ins_length $INSIZE -ins_length_sd $INSIZESD"
vopts="$iopts -max_gap_count 5 " 
oopts="-scaffolding yes -min_pair_count 3 -edgeFractionCutoff 0.03 -min_trans_lgth 200 $iopts"

echo "#START `date` " 
cd $datad
mkdir $outdir
cd $outdir

#..... run loop for vel kmer steps
shopt -s nullglob

kseqdir=vel${dv}_seq
if [ ! -f $kseqdir/Sequences ]; then
 $velbin/velveth $kseqdir 27 $vopth -noHash -shortPaired -fmtAuto -separate $inpe
fi

for k in $kset;  do { 
  ksubdir=vel${dv}_$k
  # for reruns with partial results..
  if [ -f $ksubdir/contigs.fa ]; then continue; fi
  echo "#.. start velrun $ksubdir : `date`"
  velbin=$velbin4; 
  if [ $k -le 99 ]; then velbin=$velbin2; fi
  if [ $k -le 61 ]; then velbin=$velbin1; fi
  mkdir $ksubdir
  ln -s ../$kseqdir/Sequences $ksubdir/
  $velbin/velveth $ksubdir $k $vopth -reuse_Sequences
  $velbin/velvetg $ksubdir $vopts -read_trkg yes 

  echo "#.. end velvetg $ksubdir : `date`"
} done
## end loop 1

## loop 2 oases, 1cpu; save memory, oases doesnt use multicpu
export OMP_NUM_THREADS=2
export OMP_THREAD_LIMIT=2
i=0; 
for k in $kset;  do {
  ksubdir=vel${dv}_$k
  if [ -f $ksubdir/transcripts.fa ]; then continue; fi
  if [ -f $ksubdir/contigs.fa ]; then
    echo "#.. start oases $ksubdir : `date`"
    ## DONT fork here save mem ..
    velbin=$velbin4;
    if [ $k -le 99 ]; then velbin=$velbin2; fi
    if [ $k -le 61 ]; then velbin=$velbin1; fi
    $velbin/oases   $ksubdir $oopts 
    i=$(( $i + 1 ))
  fi
} done

wait

## add cleanups, where transcripts.fa  is ok .. all Graph files?
## gzip transcripts,contigs,other keepers?
for k in $kset;  do {
  ksubdir=vel${dv}_$k
  if [ -f $ksubdir/transcripts.fa ]; then
    /bin/rm $ksubdir/{Graph2,LastGraph,PreGraph,Roadmaps}
    gzip --fast $ksubdir/{contig-ordering.txt,contigs.fa,stats.txt}
    # gzip $ksubdir/transcripts.fa later
  fi
} done


echo "#DONE : `date`"
```
