## Supplemental Archive 4 for "Genes of the Pig, *Sus scrofa*, reconstructed with EvidentialGene": srun_comet.sh

```
#! /bin/bash
## env trset=mygenes.tr datad=`pwd` prog=./runtr2cds.sh sbatch srun_prog.sh
## comet.sdsc sbatch update; show_accounts ?? not ind114
#SBATCH -A ind114
#SBATCH --job-name="tr2cds"
#SBATCH --output="tr2cds.%j.%N.out"
#SBATCH --partition=compute
#SBATCH --nodes=1
#SBATCH --ntasks-per-node=24
#SBATCH -t 39:55:00
#SBATCH --export=ALL

## 2/20cpu blastn failed evg2, try ncpu=16
export ncpuall=24
export maxmem=120000

if [ "X" = "X$datad" ]; then echo "ERR:datad=what?"; exit -1; fi
if [ "X" = "X$ncpu" ]; then ncpu=16; fi
if [ "X" = "X$prog" ]; then echo "ERR:prog=what?"; exit -1; fi
cd $datad
if [ "X" != "X$name" ]; then echo "#runname=$name datad=$datad"; fi

## FIXME: cd-hit at least is OMP threaded app, add the ENV for it, use mrun_comet.sh??
export OMP_NUM_THREADS=$ncpu

## * this is wacky like aprun/ bigred in that it calls $prog ncpu times !! need opt to not do that
echo ibrun --npernode 1 -v $prog
ibrun --npernode 1 -v $prog
#---------
```
